# The builder’s legacy: persistent reef structure creates a window for coral recovery after adult loss

**DOI:** 10.64898/2026.09.11.751047

**Authors:** Mollie Asbury, Courtney Couch, Thomas Oliver, Jessica Reichert, Joshua S. Madin

## Abstract

Coral reef recovery depends on successful juvenile recruitment, yet which local factors govern early coral demography remains poorly resolved. Adult coral cover and reef structural complexity are widely hypothesized to regulate juvenile success, but their independent and interactive effects have not been mechanistically disentangled, nor has the relative role of living adult tissue versus persisting structural framework been clarified. Here, combining *in situ* coral size-structure surveys with high resolution photogrammetry across 405 reef segments spanning the main Hawaiian Islands, we quantify how adult cover and three-dimensional reef structure, measured as rugosity and fractal dimension, jointly shape juvenile coral densities. We reveal nonlinear relationships between juvenile density and adult cover, peaking at intermediate (~8%) coral cover, with fine-scale surface complexity and vertical relief exerting opposing influences across the cover gradient. Together, these reveal a structural legacy window in which reduced adult cover relaxes constraints on juvenile establishment while the three-dimensional habitat built by those adults persists. This decoupling of biological loss from structural loss creates a transient opportunity for population renewal, with intermediate-cover reefs supporting juveniles across the broadest range of structural configurations. More generally, our results show how the physical legacy of ecosystem engineers can continue to regulate population recovery after the organisms that create it have declined.

## Introduction

All populations have environmental and habitat contexts under which growth and replenishment are maximized (Matthiopoulos et al. 2015). Across many ecological systems, population performance reflects tradeoffs between opposing constraints, for example resource limitation at one extreme and density-dependent competition at the other (MacArthur 1970, Holling 1973, Watkinson 1985, Putman et al. 1996, Badgley & Fox 2008). For reef building corals, population trajectories emerge from the balance between recruitment, survival, and growth across life stages (Connell 1973, Hughes & Tanner 2000), and demographic outcomes are particularly sensitive to processes operating during recruitment and early post-settlement growth (Hughes 1996, Doropoulos et al. 2015). Because corals are the primary architects of reef habitat, the successful transition of juveniles to reproductive adults is the fundamental demographic process underpinning reef recovery; without sustained juvenile survival and growth, adult populations cannot replenish following disturbance, and the structural complexity that supports broader reef biodiversity and function erodes over time (Hughes & Tanner 2000, Torres-Pulliza et al. 2020). Because survival and growth differ across juvenile size classes, size-structured surveys can track both immediate post-settlement survival (0–1 cm) and longer-term early persistence (1.1–5 cm; Doropoulos et al. 2015, Koester et al. 2021), providing a demographic framework for understanding recovery potential.

While broad-scale environmental and oceanographic factors, particularly larval supply, are well-established drivers of juvenile coral dynamics (Hughes et al. 2000; Hughes et al. 2002, Couch et al. 2023, Doropoulos et al. 2025), the finer-scale processes that determine whether juveniles survive following settlement remain less understood, particularly under *in situ* conditions on natural reef habitat (Pineda et al. 2010). At local scales, two mechanisms are primarily proposed to drive juvenile coral densities: adult coral cover and reef structural complexity. Adult corals play a dual role: they are both the source of new individuals through larval production and fragmentation (Vermeij 2005, Pedersen et al. 2019) and the architects of the three-dimensional structural framework that shapes reef habitat. Structural complexity provides microhabitats that increase settlement by reducing juvenile mortality and shaping the fine-scale hydrodynamic conditions that govern larval settlement and survival (Yadav et al. 2016, Reidenbach et al. 2021, Carlson et al. 2024). Because reef structure is built by, but not exclusive to, living adult coral tissue, these two roles can be decoupled: the physical architecture of the reef may persist and function as juvenile habitat independently of the living adults that originally produced it. Whether it is the biological presence of adults or the structural complexity they generate that more directly governs juvenile success has not been fully resolved, yet the distinction has important consequences for understanding what reef conditions promote recovery.

The relative influence of adult cover and structural complexity likely shift following settlement, as recently settled recruits experience the highest mortality immediately and are strongly influenced by fine-scale microhabitat availability, while larger juveniles are increasingly constrained by competition for space and resources (Ritson-Williams et al. 2009, Penin et al. 2010). Disentangling these processes therefore requires not only resolving how adult cover and structure interact, but also how that interaction varies across early ontogeny. Here, we address this by combining *in situ* coral size-structure surveys with high-resolution photogrammetry across 405 reef segments spanning the main Hawaiian Islands. We use complementary metrics of three-dimensional reef structure—rugosity and fractal dimension—that independently capture surface area and niche breadth (Torres-Pulliza et al. 2020, Schiettekatte et al. 2025), allowing their interactive effects with adult cover to be resolved across post-settlement and established juvenile size classes. The main Hawaiian Islands provide a compelling study system, with relatively steady larval supply and recruitment rates at the upper end of ranges observed across the Pacific (Price et al. 2019), even after severe heat stress (Couch et al. 2023). Variation in reef habitat types across the region enables assessment of broad gradients in structural complexity and coral cover (Asbury et al. 2023), yet juvenile densities vary widely within and among islands in ways not consistently explained by environmental conditions alone (Brown 2004, Coles & Brown 2007). Our approach focuses on the response side of recruitment, specifically the post-settlement bottlenecks that determine whether reefs can recover once larvae arrive, and d by doing so, tests how the biological presence of adults and the structural complexity they create interact to determine where juvenile corals can establish, and whether their decoupling creates conditions that favor population recovery.

## Results

Study sites across the main Hawaiian Islands spanned broad gradients in juvenile coral density, adult coral cover, rugosity, and fractal dimension (Figure 1). Juvenile densities varied widely among sites, with most sites characterized by low densities and a smaller number exhibiting very high juvenile densities (Figure 1B). Adult coral cover similarly spanned low to high values across the region (Figure 1C). Measures of habitat structure showed substantial variability, with rugosity and fractal dimension differing both within and among islands (Figure 1D, E).

**Figure 1.**
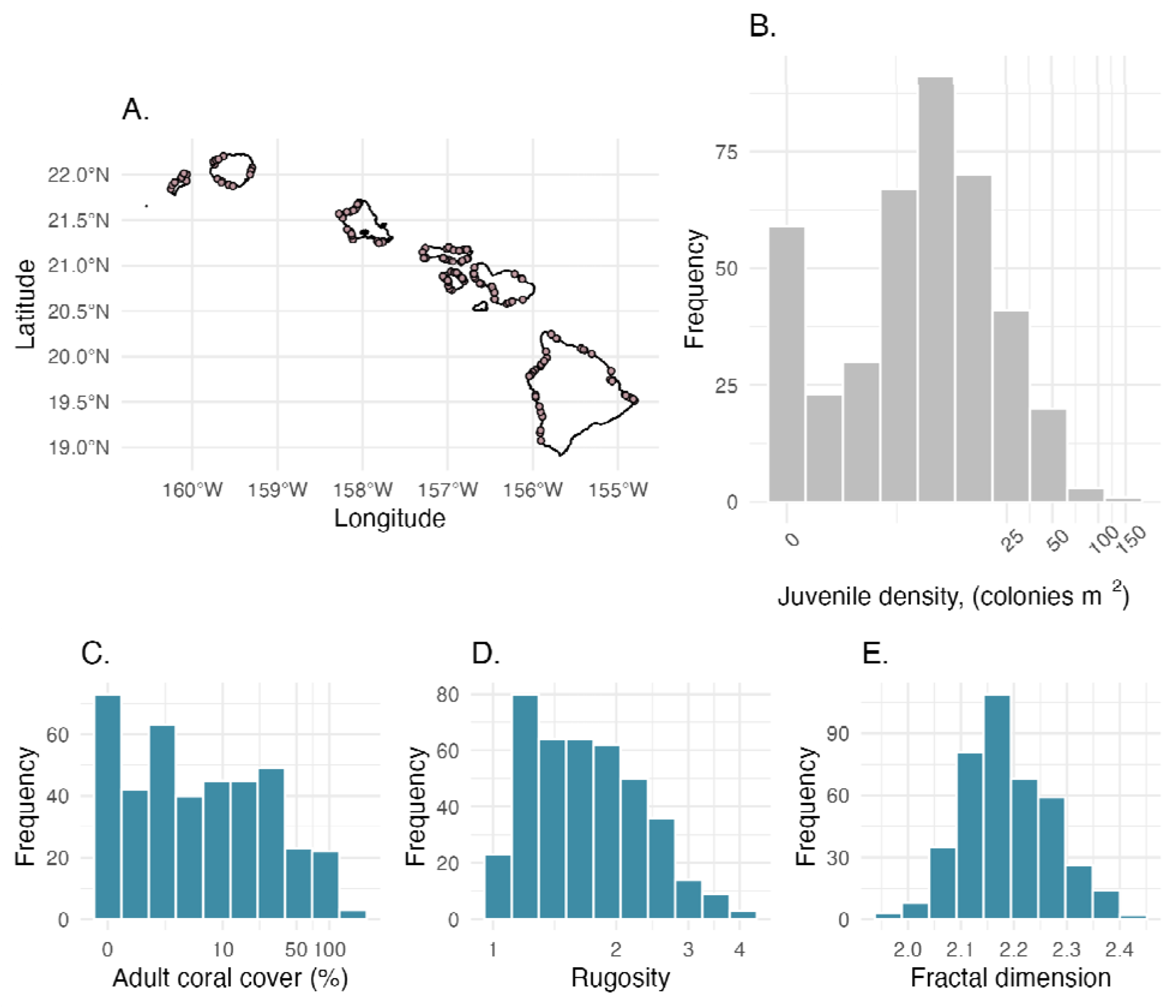
(A) Map of 143 survey sites in the Hawaiian Archipelago. (B) Frequency distribution of total juvenile coral density. (C-E) Distributions of adult coral cover, rugosity, and fractal dimension.

### The role of adult coral cover and reef structure

Total juvenile density was best predicted by a nonlinear effect of adult coral cover, the interaction between coral cover and fractal dimension, and the interaction between rugosity and fractal dimension, in order of effect size (Table S2, Figure S1A; pseudo-R_2_ = 0.72 [Cragg-Uhler], 0.11 [McFadden]). Juvenile density showed a strong unimodal relationship with adult cover, peaking at approximately 8% under mean structural conditions, but the position of this peak depended strongly on reef structure (Figure S1B). Fractal dimension modulated the effect of coral cover on juvenile density: at low fractal dimension, predicted juvenile density peaked at around 4% adult coral cover, whereas at high fractal dimension the peak shifted to approximately 14% (Figure S1C). The interaction between rugosity and fractal dimension further revealed that the effect of rugosity depended on the level of fractal dimension. At high fractal dimension, rugosity showed a negative relationship with juvenile density, while at low fractal dimension increasing rugosity predicted higher juvenile density; at intermediate fractal dimension, rugosity had little effect (Figure S1D).

### Divergent size class responses

Established juvenile density (1.1–5 cm) was three-fold higher than post-settlement density (0–1 cm; Figure 2A). The two size classes showed distinct relationships with adult coral cover and reef structure. Post-settlement juvenile density was most strongly influenced by the interaction between coral cover and rugosity, the interaction between coral cover and fractal dimension, the linear effect of adult coral cover, and the interaction between rugosity and fractal dimension, in order of effect (Table S2; Figure 2B; pseudo-R_2_ = 0.41 [Cragg-Uhler], 0.12 [McFadden]). Post-settlement density decreased with increasing coral cover (Figure 2C), and the magnitude of this negative relationship depended on reef structure. Higher rugosity slightly intensified the negative effect of coral cover while higher fractal dimension lessened it (Figure S2A, B). Established juvenile density was most strongly influenced by a nonlinear effect of coral cover and, to a lesser extent, the interactions between coral cover and fractal dimension and between rugosity and fractal dimension (Table S2; Figure 2B; Figure S2C; expanded on below; pseudo-R_2_ = 0.59 [Cragg-Uhler], 0.10 [McFadden]). In contrast to post-settlement juveniles, established juveniles showed a strong unimodal relationship with coral cover, with density peaking at approximately 8% under mean structural conditions (Figure 2C). Thus, the relative importance of structure and adult cover shifted across early ontogeny: post-settlement densities were most strongly associated with habitat configuration, whereas densities of juveniles that had persisted beyond the earliest life stage showed the strongest constraint from increasing adult cover.

**Figure 2.**
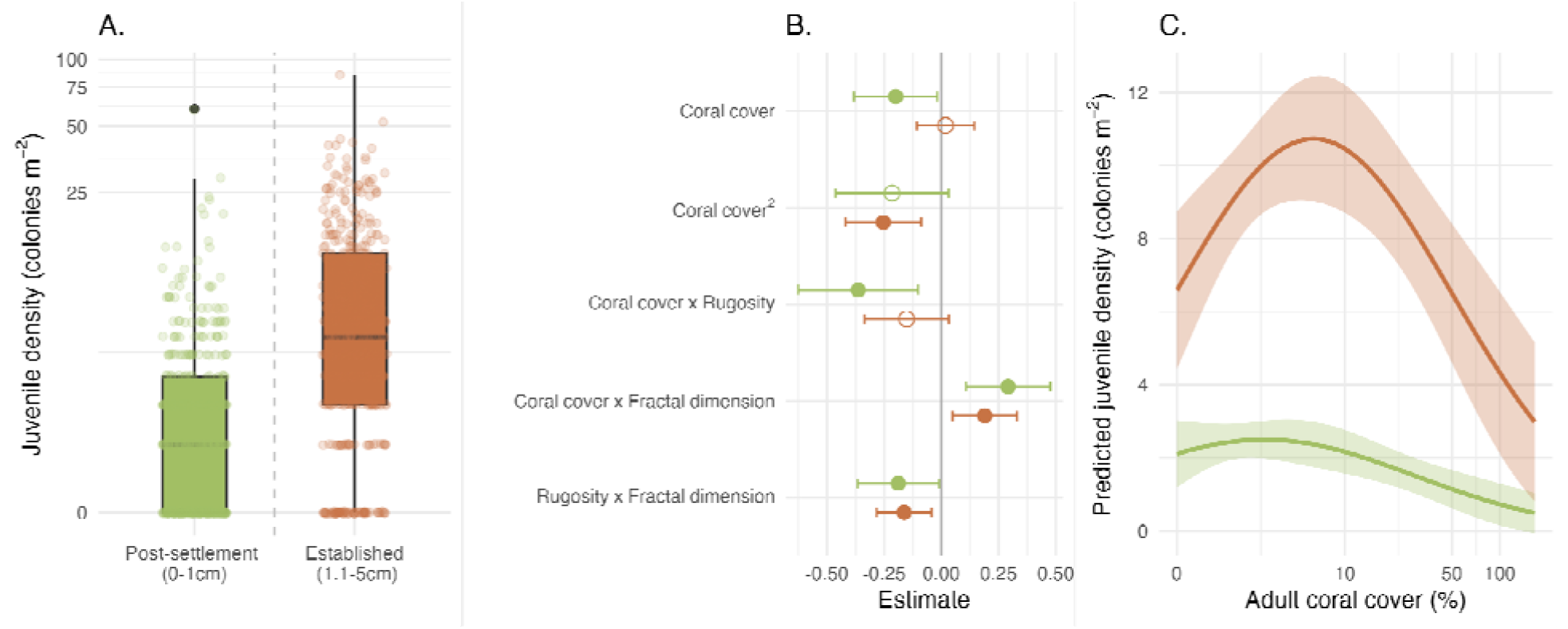
The effects of adult coral cover and reef structural complexity on two size classes of juvenile corals in the Hawaiian Archipelago: post-settlement (0–1 cm; green) and established (1.1–5 cm; orange). (A) Distribution of juvenile density classified as post-settlement (0–1 cm) and established (1.1–5 cm) size classes. (B) Parameter estimates from survey-weighted Poisson regression predicting density of post-settlement juveniles and established juveniles. Error bars represent the 95% confidence interval and color corresponds to size class. Filled points indicate that confidence intervals do not overlap with zero and are therefore considered significant. (C) Predicted post-settlement and established juvenile density as a function of adult coral cover. Shaded ribbons represent 95% confidence intervals and color corresponds to size class. For predictions, all other covariates were held at their mean. Plotted trends of other significant covariates are depicted in Figure S2.

### Intermediate coral cover broadens structural suitability

When predicted densities were examined across the full range of rugosity and fractal dimension at fixed coral cover levels, the structural configurations supporting juvenile corals shifted markedly with coral cover and the breadth of favorable conditions varied among cover levels (Figure S3). The juvenile suitability index, integrating scaled predictions for both size classes simultaneously, captured these patterns and revealed their cover-dependence most clearly (Figure 3). At high adult coral cover (75%), suitability hotspots for both size classes were concentrated at high fractal dimension and low rugosity (Figure 3A), indicating a narrow range of structural conditions supporting juveniles. At intermediate cover (8%), two distinct suitability peaks emerged—one at high rugosity and low fractal dimension, and a second at high fractal dimension and low rugosity—reflecting a substantially broader range of structural configurations that support juvenile success than at either high or low cover (Figure 3B). At low cover (1%), suitability hotspots shifted back toward high rugosity and low fractal dimension, with a narrower range of favorable configurations than at intermediate cover (Figure 3C). This pattern was quantified directly as habitat breadth, calculated as the percentage of combinations of rugosity and fractal dimension with joint suitability exceeding 0.50, which was highest at intermediate cover (6.8%) relative to high (1.1%) and low (2.6%) cover (Figure 3D). Thus, intermediate-cover reefs were distinguished not only by high predicted juvenile densities, but by a substantially broader range of reef structures capable of simultaneously supporting both juvenile size classes.

**Figure 3.**
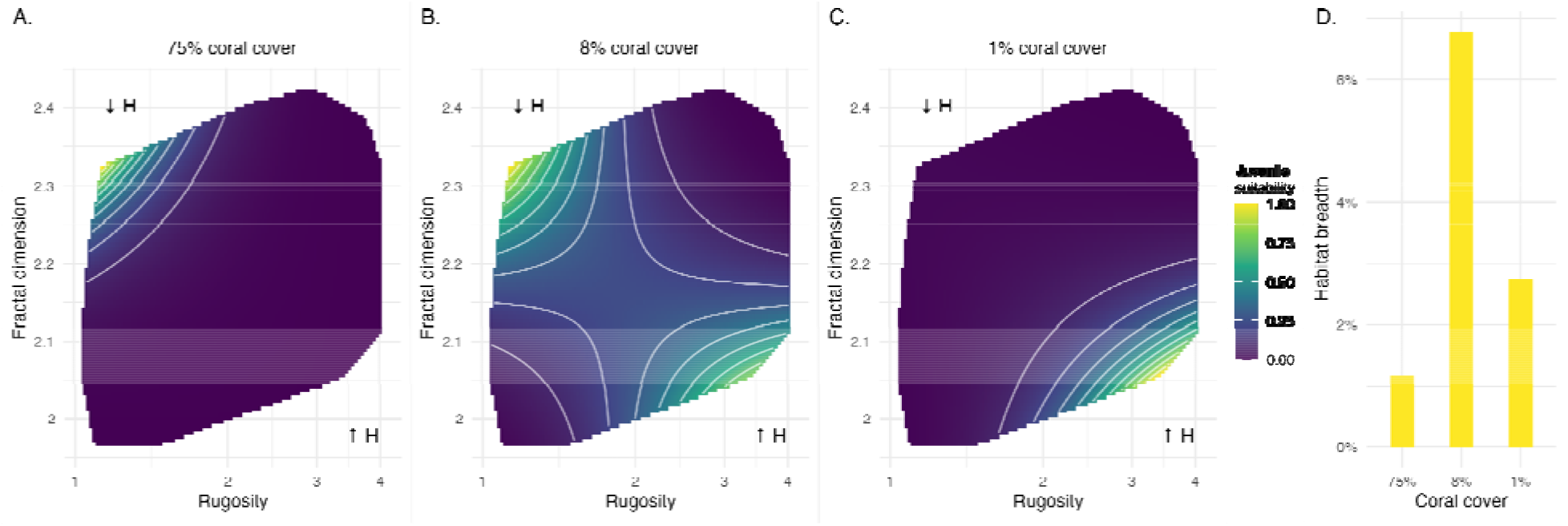
Structural configurations and juvenile habitat suitability across coral cover states. (A-C) Juvenile suitability index across combinations of rugosity and fractal dimension at 75%, 8%, and 1% adult coral cover. The index integrates scaled predicted densities for post-settlement and established juveniles simultaneously, ranging from 0 (low suitability) to 1 (high suitability). Arrows denote high ( H) and low ( H) height ranges according to the geometric relationship between rugosity and fractal dimension. (D) Habitat breadth at each coral cover level, defined as the percentage of rugosity x fractal dimension combinations with joint post-settlement and post-established juvenile suitability exceeding 0.50.

## Discussion

Understanding coral reef recovery requires not only identifying correlates of juvenile recruitment but resolving the mechanisms that generate them. Our results reveal a fundamental paradox in reef recovery: the conditions that maximize adult coral cover are not those that maximize juvenile success. Juveniles thrive in the more open, more diverse habitat provided by intermediate levels of cover. Adult corals simultaneously create the three-dimensional habitat used by the next generation and constrain juvenile establishment through space occupation and competition. Crucially, these opposing effects operate on different timescales: adult mortality can rapidly release biological constraints, while the physical structure created by those adults persists. We propose that this temporal decoupling creates a structural legacy window in which recruitment habitat remains after much of the living coral that created it has disappeared. The result is not simply higher juvenile density at intermediate cover, but a broader range of structural configurations capable of supporting juvenile corals. Structural legacy therefore provides a mechanism by which the decline of an ecosystem engineer can temporarily create conditions favorable for its own replacement.

The peak in juvenile density at intermediate adult cover is consistent with patterns reported elsewhere (~10-12%; Hughes 1985, Vermeij & Sandin 2008, Couch et al. 2023), suggesting that our unimodal relationship itself may be widespread across reef systems. Our results provide a potential mechanistic explanation for this pattern: adult cover and the habitat created by adults become partially decoupled, while fine-scale structural configuration determines the range of conditions under which juveniles can exploit this release from adult competition. Critically, structural complexity does not universally benefit juvenile corals, but configuration matters more than magnitude. High vertical relief combined with tightly packed micro-crevices, the combination not associated with higher juvenile suitability (Figure 3), likely reduces juvenile persistence by trapping sediments or promoting algal accumulation inaccessible to herbivores (Brandl et al. 2014). Structural complexity expands niche availability and relaxes density-dependent constraints (Kovalenko et al. 2012, Edmunds et al. 2018), but only when spatial configuration produces accessible refugia. Coral cover alone is therefore an incomplete proxy for recovery potential. The structural configuration of available habitat matters as much as its quantity, and neither variable can substitute for the other.

The distinct responses of the two juvenile size classes reveal how the dominant demographic bottleneck shifts across early ontogeny. Post-settlement juveniles (0–1 cm) were most sensitive to fine-scale habitat configuration, declining with increasing coral cover but buffered by higher fractal dimension, consistent with micro-habitat driven responses at the settlement stage (Penin et al. 2010, Carlson et al. 2024). Established juveniles (1.1–5 cm) were more strongly governed by adult coral cover, exhibiting a unimodal relationship that peaks at approximately 8%, consistent with space limitation and density-mediated processes becoming the dominant constraint as juveniles grow (Connell 1978, Hughes 1985). Rather than a categorical shift, these patterns reflect a gradual transition in which structural complexity governs the earliest bottleneck, whether a settler survives its first weeks, while adult cover governs the subsequent bottleneck, whether an established juvenile can persist in an increasingly crowded assemblage.

Resolving both bottlenecks simultaneously requires conditions where structure is accessible and cover is moderate: precisely the structural legacy window. The cover-dependent patterns in juvenile suitability across the rugosity-fractal dimension surface clarify what makes intermediate cover reefs ecologically distinctive and suggest a three-phase hypothesis for how juvenile habitat may change through disturbance and recovery. In the first undisturbed phase, with high competition at 75% coral cover, living adult corals occupy space and build a three-dimensional structural framework, but juvenile success is constrained to a narrow range of structural configurations where complex surface geometry at low vertical relief buffers competitive exclusion. In phase 2 (~8% coral cover), what we term the structural legacy window, partial adult mortality reduces competitive pressure while structural complexity remains elevated despite declining live cover, decoupling reef complexity from adult cover and creating the conditions of the broadest juvenile habitat suitability, more than double both other phases. Our data are consistent with this state: moderate coral cover reefs driving the juvenile suitability peak represent systems where competition has been partially released without degrading the structural framework that facilitates juvenile success. In the final phase of successional transition with very little coral cover, competition is relaxed and absolute juvenile densities can be high, but only under a structurally specific combination of high vertical relief and relatively simple surface geometry, a configuration that may reflect legacy structural framework from previously dense coral communities (Alvarez-Filip et al. 2009, Alvarez-Filip et al. 2013) and one that becomes increasingly rare as bioerosion and physical degradation erode that framework over time (Perry et al. 2013, Cornwall et al. 2026). This structural permissiveness, or the capacity to support juveniles regardless of the specific structural state of the reef, is what distinguishes intermediate cover reefs as recovery opportunities. Therefore, recovery does not depend on achieving a particular structural configuration; it is robust to the variability that characterizes partially disturbed reefscapes. Our spatial data therefore generate a testable temporal prediction: following partial coral mortality, juvenile establishment should initially increase as adult constraints are released while structural habitat persists, before declining as that structural legacy erodes. Confirming this predicted sequence will require temporal monitoring of individual reefs through disturbance and recovery.

These findings challenge a central assumption of reef resilience assessment: that live coral cover is the primary indicator of recovery potential. Our results suggest that structural persistence, or the three-dimensional complexity that outlasts living tissue, is an equally important and currently under-monitored determinant of where juvenile success is possible. A reef assessed as degraded by live cover metrics may retain broad recovery potential if its structural framework remains intact and moderate cover is maintained. Conversely, a reef at high live cover where structure has been simplified may offer fewer viable microhabitats for juvenile persistence than its cover alone would suggest. Incorporating structural complexity alongside live cover into resilience frameworks would better capture the true demographic state of reefs across their disturbance trajectories. Reef structure is amenable to local intervention through protection of surviving structural frameworks, restoration of lost habitat complexity, and management actions that extend the period over which structurally intact reefs remain capable of natural replenishment. Until structural persistence is measured alongside live cover, reef resilience assessments will continue to misidentify the reefs most capable of recovery. Understanding the relative rates at which living cover and engineered structure are lost is therefore critical for predicting whether disturbed reefs retain the capacity for natural population recovery, thereby translating reef resilience theory into practice.

## Methods

### Study design and field surveys

Surveys were conducted between April and July 2019 as part of NOAA’s National Coral Reef Monitoring Program (NCRMP) across seven islands of the main Hawaiian Islands: Hawai’i, Maui, Lāna□i, Moloka□i, O□ahu, Kaua□i, and Ni□ihau (Figure 1A). Sites were selected following stratified random sampling design within hardbottom forereef habitats stratified into three depth bins (shallow: 0-6m, mid: >6-18m, deep: >18-30m), with sampling effort proportional to hardbottom area and variance in coral density from prior years (Smith et al. 2011). At each site, a 30m transect line was deployed parallel to the depth contour. Coral demographic surveys were conducted within three to four 1 x 2.5m segments spaced evenly along the transect (Winston et al. 2019). For adult corals (>5 cm maximum diameter) whose center of mass fell within the segment, divers recorded lowest taxonomic identification, maximum diameter, and morphology. Juvenile corals (0-5 cm) were recorded within the first 1×1m of each segment, measured, and identified to genus level.

### Photogrammetry and structural complexity

A Structure-from-Motion (SfM) photogrammetry survey was conducted at each site prior to coral demographic surveys using a Canon EOS Rebel SL2 digital camera, covering 50-80 m_2_ with three to four ground control points. High-resolution reef models were constructed in Agisoft Metashape following standard protocol (Suka et al. 2019, Torres-Pulliza et al. 2024), and digital elevation models (DEMs) were extracted at 0.002m resolution. Segments were delineated in QGIS using orthomosaics and segment shapefiles were used to crop site-level DEMS, with NA values filled by inverse distance weighting. Two complementary metrics of structural complexity were estimated in R using the habtools package (Schiettekatte et al. 2025): rugosity, calculated using the area method at 1 cm resolution; and fractal dimension estimated using the theory method at 1 cm resolution. These metrics were chosen for their geometric interdependence, which allows patterns in rugosity and fractal dimension to inform inferences about height range (Torres-Pulliza et al. 2020).

### Coral cover and juvenile density estimates

Segment-level adult coral cover was derived from diameter–area relationships for different species and morphologies, with coefficients estimated from 4,718 digitally outlined colonies spanning multiple islands (Table S1). For species or morphology combinations not represented in the dataset, coefficients from the most geometrically similar category were used. Partial mortality was excluded from colony area. Adult coral cover was calculated as summed colony area divided by segment area (2.5 m^2^); segments with cover >100% (n = 10) were retained as these may reflect true colony overlap. Adult coral cover was additionally split by genus, morphology, and life-history strategy (LHS; Darling et al. 2012) to test more descriptive estimations of the adult population; given the depauperate morphological assemblage of the main Hawaiian Islands, comprising primarily encrusting, mounding, and branching forms with no tables or arborescent species, the functional variance captured by these splits is limited relative to more diverse Indo-Pacific assemblages, and models incorporating these splits showed no significant effects and were excluded from final analyses. Juvenile coral density was summarized for total (0–5 cm), post-settlement (0–1 cm), and established (1.1–5 cm) size classes, corresponding to early post-settlement recruits and juveniles that have survived initial mortality (Ritson-Williams et al. 2009; Penin et al. 2010).

### Statistical analysis

We used the survey package in R (Lumley 2024) for all models, with inverse probability weights based on selection probability within strata and a nested structure defined by island, sector, and depth bin. Juvenile coral density was modelled as a function of adult coral cover and structural complexity using survey-weighted generalized linear models with a Poisson distribution and log link (svyglm). Adult coral cover and rugosity were log-transformed; all predictors were scaled and centered. The full model included second-order polynomial terms for adult coral cover and pairwise interactions between coral cover, rugosity, and fractal dimension. Separate models were fitted for total juvenile density, post-settlement density, and established juvenile density. Predictions were generated across a grid of coral cover values with rugosity and fractal dimension held at their mean to visualize marginal effects, and across combinations of rugosity and fractal dimension at three ecologically representative cover levels—1%, 8%, and 75%—capturing the low, intermediate, and high regions of the response surface. To evaluate conditions simultaneously supporting both size classes, a juvenile suitability index was calculated for each cover level as the product of scaled (0–1) predicted densities from post-settlement and established juveniles.

## Acknowledgements

We would like to thank the crew of the NOAA vessel Oscar Elton Sette for providing field support during the main Hawaiian Islands National Coral Reef Monitoring Programs cruise. We would also like to thank all participating scientists for collection of Structure-from-Motion imagery and underwater coral surveys. This study was funded by the Defense Advanced Research Projects Agency [BAA HR001121S0012 to M.A., J.R., and J.S.M.], the Directorate for Geosciences [1948946 to M.A., J.R., and J.S.M.], and the National Oceanic and Atmospheric Administration’s Coral Reef Conservation Program [Project #743 to M.A., C.C., and T.O.]. The funders played no role in study design, analysis and interpretation of data, or the writing of this manuscript.

## Data availability

The data and underlying code supporting the findings of this study are available in Figshare via https://figshare.com/s/8e9326a26c22201ff70c. Upon acceptance of the manuscript, the code will be permanently archived and made publicly available on Github.

## Notes

### Competing Interest Statement

The authors have declared no competing interest.

## References

Alvarez-Filip, L., Dulvy, N. K., Gill, J. A., Côté, I. M. & Watkinson, A. R. (2009). Flattening of Caribbean coral reefs: region-wide declines in architectural complexity. Proc. R. Soc. Lond., Ser. B: Biol. Sci. 276, 3019–3025.

Alvarez-Filip, L., Carricart-Ganivet, J., Horta-Puga, G. et al. (2013). Shifts in coral-assemblage composition do not ensure persistence of reef functionality. Sci Rep 3, 3486. 10.1038/srep03486

Asbury, M., Schiettekatte, N. M., Couch, C. S., Oliver, T., Burns, J. H., & Madin, J. S. (2023). Geological age and environments shape reef habitat structure. Global Ecology and Biogeography, 32(7), 1230–1240.

Badgley, C., & Fox, D. L. (2000). Ecological biogeography of North American mammals: species density and ecological structure in relation to environmental gradients. Journal of Biogeography, 27(6), 1437–1467.

Brandl, S. J., Hoey, A. S., & Bellwood, D. R. (2014). Micro-topography mediates interactions between corals, algae, and herbivorous fishes on coral reefs. Coral Reefs, 33(2), 421–430.

Brown, E. K. (2004) Reef coral populations: Spatial and temporal differences observed at six reefs off West Maui. Ph.D. Thesis, University of Hawaii, p 277.

Carlson, R. R., Crowder, L. B., Martin, R. E., & Asner, G. P. (2024). The effect of reef morphology on coral recruitment at multiple spatial scales. Proceedings of the National Academy of Sciences, 121(4), e2311661121.

Coles, S. L., & Brown, E. K. (2007). Twenty-five years of change in coral coverage on a hurricane impacted reef in Hawai ‘i: the importance of recruitment. Coral Reefs, 26(3), 705–717.

Connell, J. H. (1973). Population ecology of reef-building corals. Biology and Geology of Coral Reefs, 2, 205–245.

Connell, J. H. (1978). Diversity in tropical rain forests and coral reefs: high diversity of trees and corals is maintained only in a nonequilibrium state. Science, 199(4335), 1302–1310.

Cornwall, C.E., Timmerman, O., Andersson, A. et al. (2026). Persistence of coral reef structures into the twenty-first century. Nat Rev Earth Environ 7, 151–161. 10.1038/s43017-026-00764-4

Couch, C. S., Oliver, T. A., Dettloff, K., Huntington, B., Tanaka, K. R., & Vargas-Ángel, B. (2023). Ecological and environmental predictors of juvenile coral density across the central and western Pacific. Frontiers in Marine Science, 10, 1192102.

Darling, E. S., Alvarez-Filip, L., Oliver, T. A., McClanahan, T. R., & Côté, I. M. (2012). Evaluating life-history strategies of reef corals from species traits. Ecology Letters, 15(12), 1378–1386.

Doropoulos, C., Ward, S., Roff, G., González-Rivero, M., & Mumby, P. J. (2015). Linking demographic processes of juvenile corals to benthic recovery trajectories in two common reef habitats. PLoS One, 10(5), e0128535.

Doropoulos, C., Alvarez-Noriega, M., Fabricius, K., Ferrari, R., Mumby, P. J., Noonan, S. H., … & Salee, K. (2025). Impact of environmental gradients on juvenile coral demography across the Great Barrier Reef and Torres Strait. Coral Reefs, 1–17.

Edmunds, P. J., Nelson, H. R., & Bramanti, L. (2018). Density-dependence mediates coral assemblage structure. Ecology, 99(11), 2605–2613.

Holling, C. S. (1973, November). Resilience and stability of ecological systems.

Hughes, T. P. (1985). Life histories and population dynamics of early successional corals. Antenne Museum-EPHE.

Hughes, T. P. (1996). Demographic approaches to community dynamics: a coral reef example. Ecology, 77(7), 2256–2260.

Hughes, T. P., Baird, A. H., Dinsdale, E. A., Moltschaniwskyj, N. A., Pratchett, M. S., Tanner, J. E., & Willis, B. L. (2000). Supply-side ecology works both ways: The link between benthic adults, fecundity, and larval recruits. Ecology, 81(8), 2241–2249.

Hughes, T. P., Baird, A. H., Dinsdale, E. A., Harriott, V. J., Moltschaniwskyj, N. A., Pratchett, M. S., … & Willis, B. L. (2002). Detecting regional variation using meta-analysis and large-scale sampling: latitudinal patterns in recruitment. Ecology, 83(2), 436–451.

Hughes, T. P., & Tanner, J. E. (2000). Recruitment failure, life histories, and long-term decline of Caribbean corals. Ecology, 81(8), 2250–2263.

Koester, A., Ford, A. K., Ferse, S. C., Migani, V., Bunbury, N., Sanchez, C., & Wild, C. (2021). First insights into coral recruit and juvenile abundances at remote Aldabra Atoll, Seychelles. PLoS One, 16(12), e0260516.

Kovalenko, K. E., Thomaz, S. M., & Warfe, D. M. (2012). Habitat complexity: approaches and future directions. Hydrobiologia, 685(1), 1–17.

Lumley, T. (2024). “survey: analysis of complex survey samples.” R package version 4.4.

MacArthur, R. (1970). Species packing and competitive equilibrium for many species. Theoretical population biology, 1(1), 1–11.

Matthiopoulos, J., Fieberg, J., Aarts, G., Beyer, H. L., Morales, J. M., & Haydon, D. T. (2015). Establishing the link between habitat selection and animal population dynamics. Ecological Monographs, 85(3), 413–436.

Pedersen, N. E., Edwards, C. B., Eynaud, Y., Gleason, A. C., Smith, J. E., & Sandin, S. A. (2019). The influence of habitat and adults on the spatial distribution of juvenile corals. Ecography, 42(10), 1703–1713.

Penin, L., Michonneau, F., Baird, A. H., Connolly, S. R., Pratchett, M. S., Kayal, M., & Adjeroud, M. (2010). Early post-settlement mortality and the structure of coral assemblages. Marine Ecology Progress Series, 408, 55–64.

Perry, C., Murphy, G., Kench, P. et al. Caribbean-wide decline in carbonate production threatens coral reef growth. Nat Commun 4, 1402 (2013). 10.1038/ncomms2409

Pineda, J., Porri, F., Starczak, V., & Blythe, J. (2010). Causes of decoupling between larval supply and settlement and consequences for understanding recruitment and population connectivity. Journal of Experimental Marine Biology and Ecology, 392(1-2), 9–21.

Price, N. N., Muko, S., Legendre, L., Steneck, R., van Oppen, M. J., Albright, R., … & Edmunds, P. J. (2019). Global biogeography of coral recruitment: tropical decline and subtropical increase. Marine Ecology Progress Series, 621, 1–17.

Putman, R. J., Langbein, J., M. Hewison, A. J., & Sharma, S. K. (1996). Relative roles of density-dependent and density-independent factors in population dynamics of British deer. Mammal Review, 26(2-3), 81–101.

Reidenbach, M. A., Stocking, J. B., Szczyrba, L., & Wendelken, C. (2021). Hydrodynamic interactions with coral topography and its impact on larval settlement. Coral Reefs, 40(2), 505–519.

Ritson-Williams, R., Arnold, S. N., Fogarty, N. D., Steneck, R. S., Vermeij, M. J., & Paul, V. J. (2009). New perspectives on ecological mechanisms affecting coral recruitment on reefs. Smithsonian Contributions to the Marine Sciences, 38, 437.

Schiettekatte, N., Asbury, M., Chen, G. K., Dornelas, M., Reichert, J., Torres-Pulliza, D., Zawada, K. J. A., & Madin, J. S. (2025). habtools: An R package to calculate 3D metrics for surfaces and objects. Methods in Ecology and Evolution, 16, 895–903. 10.1111/2041-210X.70027

Smith, S. G., Swanson, D. W., Chiappone, M., Miller, S. L., & Ault, J. S. (2011). Probability sampling of stony coral populations in the Florida Keys. Environmental Monitoring and Assessment, 183(1), 121–138.

Suka, R., Asbury, M., Gray, A. E., Winston, M., Oliver, T. T. A., & Couch, C. S. (2019). Processing photomosaic imagery of coral reefs using structure-from-motion standard operating procedures.

Torres-Pulliza, D., Charendoff, J., Couch, C., Suka, R., Gray, A., Lichowski, F., … & Oliver, T. (2024). Processing coral reef imagery using Structure-from-Motion photogrammetry: Standard operating procedures (2023 update).

Torres-Pulliza, D., Dornelas, M. A., Pizarro, O., Bewley, M., Blowes, S. A., Boutros, N., … & Madin, J. S. (2020). A geometric basis for surface habitat complexity and biodiversity. Nature Ecology & Evolution, 4(11), 1495–1501.

Vermeij, M. J. A. (2005). Substrate composition and adult distribution determine recruitment patterns in a Caribbean brooding coral. Marine Ecology Progress Series, 295, 123–133.

Vermeij, M. J., & Sandin, S. A. (2008). Density-dependent settlement and mortality structure the earliest life phases of a coral population. Ecology, 89(7), 1994–2004.

Watkinson, A. R. (1985). On the abundance of plants along an environmental gradient. The Journal of Ecology, 569–578.

Winston, M., Couch, C. S., Ferguson, M., Huntington, B., Swanson, D. W., & Vargas-Ángel, B. (2019). Ecosystem sciences division standard operating procedures: Data collection for rapid ecological assessment benthic surveys, 2018 update.

Yadav, S., Rathod, P., Alcoverro, T., & Arthur, R. (2016). “Choice” and destiny: the substrate composition and mechanical stability of settlement structures can mediate coral recruit fate in post-bleached reefs. Coral Reefs, 35(1), 211–222.

