## Supplemental tables & figures for "The builder’s legacy: persistent reef structure creates a window for coral recovery after adult loss"

**Supplementary Materials**

Table S1. Diameter-area coefficients (a, b) and sample sizes (number of colonies) for each species-morphology combination in the Hawaiian Archipelago. Coefficients were derived from log-log regressions of delineated colony area against maximum diameter. Morphology only rows (“all species”) summarize coefficients across all species within that growth form.

| **Species** | **Morphology** | **a (intercept)** | **b (slope)** | **Number of Colonies** |
| --- | --- | --- | --- | --- |
| *Montipora* sp. | Encrusting | 0.686 | 1.965 | 25 |
| *Montipora capitata* | Branching | 0.785 | 2.000 | 14 |
| *Montipora capitata* | Encrusting | 0.616 | 1.936 | 1301 |
| *Montipora capitata* | Encrusting columnar | 0.076 | 1.518 | 10 |
| *Montipora capitata* | Knobby | 0.785 | 2.000 | 859 |
| *Montipora capitata* | Mounding | 0.785 | 2.000 | 8 |
| *Montipora capitata* | Plating | 0.785 | 2.000 | 10 |
| *Montipora patula* | Encrusting | 0.766 | 1.998 | 877 |
| *Montipora patula* | Encrusting columnar | 0.099 | 1.530 | 4 |
| *Montipora patula* | Knobby | 0.785 | 2.000 | 3 |
| *Montipora patula* | Mounding | 0.785 | 2.000 | 4 |
| *Montipora patula* | Plating | 0.785 | 2.000 | 2 |
| *Pocillopora meandrina* | Branching | 0.589 | 1.920 | 28 |
| *Pocillopora meandrina* | Encrusting | 0.785 | 2.000 | 6 |
| *Pocillopora meandrina* | Knobby | 0.717 | 2.001 | 31 |
| *Porites* sp. | Encrusting | 0.999 | 2.131 | 80 |
| *Porites* sp. | Mounding | 0.755 | 1.988 | 71 |
| *Porites lobata* | Encrusting | 0.290 | 1.856 | 239 |
| *Porites lobata* | Mounding | 0.338 | 1.885 | 1076 |
| *Porites lutea* | Encrusting | 0.785 | 2.000 | 23 |
| *Porites lutea* | Mounding | 0.785 | 2.000 | 43 |
| All species | Branching | 0.595 | 1.903 | 42 |
| All species | Encrusting | 0.635 | 1.959 | 2551 |
| All species | Encrusting columnar | 0.080 | 1.499 | 15 |
| All species | Knobby | 0.781 | 1.999 | 895 |
| All species | Mounding | 0.364 | 1.891 | 1202 |
| All species | Plating | 0.785 | 2.000 | 13 |

Table S2. Model results of survey-weighted Poisson regressions for (A) total juvenile density, (B) post-settlement (0–1 cm) juvenile density, and (C) established (1.1–5 cm) juvenile density. Significant variables are in bold and parentheses denote standard error. *** P < 0; ** P < 0.001; * P < 0.01; † P < 0.05

| Covariate | A. Total juvenile density | B. Post-settlement (0–1 cm) juvenile density | C. Established (1.1–5 cm) juvenile density |
| --- | --- | --- | --- |
| Coral cover | -0.03 (0.06) | **-0.21 (0.09) *** | 0.02 (0.06) |
| Coral cover ^2^ | **-0.24 (0.09) **** | -0.21 (0.13) | **-0.25 (0.08) **** |
| Coral cover * Rugosity | -0.19 (0.10) † | **-0.35 (0.13) **** | -0.15 (0.09) |
| Coral cover * Fractal dimension | **0.22 (0.07) **** | **0.30 (0.09) **** | **0.19 (0.07) **** |
| Rugosity * Fractal dimension | **-0.18 (0.06) **** | **-0.20 (0.09) *** | **-0.17 (0.06) **** |

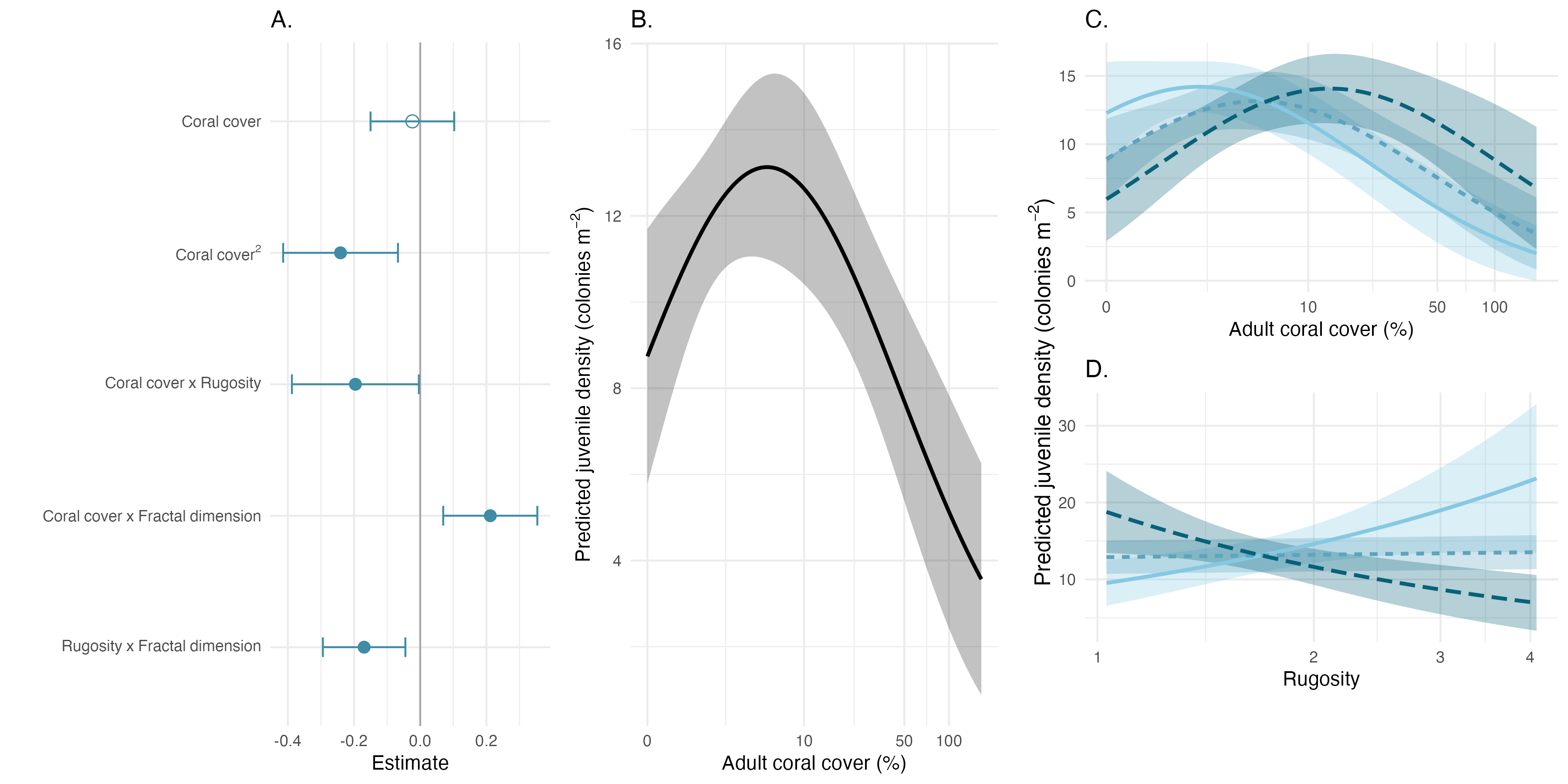

Figure S1. Predicted relationships between adult coral cover, reef structural complexity, and total juvenile coral density. (A) Parameter estimates from survey-weighted Poisson regression predicting juvenile density. Error bars represent the 95% confidence interval. Filled points indicate that confidence intervals do not overlap with zero and are therefore considered significant. (B) Predicted total juvenile density as a function of adult coral cover with 95% confidence intervals (shaded ribbon). (C) Predicted total juvenile density as a function of adult coral cover and low, medium, and high fractal dimension (D; 10th, 50th, and 90th percentiles, respectively). (D) Predicted total juvenile density as a function of rugosity and low, medium, and high fractal dimension (D; 10th, 50th, and 90th percentiles, respectively). For panels C-D, shaded ribbons represent 95% confidence intervals and line type corresponds to fractal dimension level; all other covariates were held at their mean.

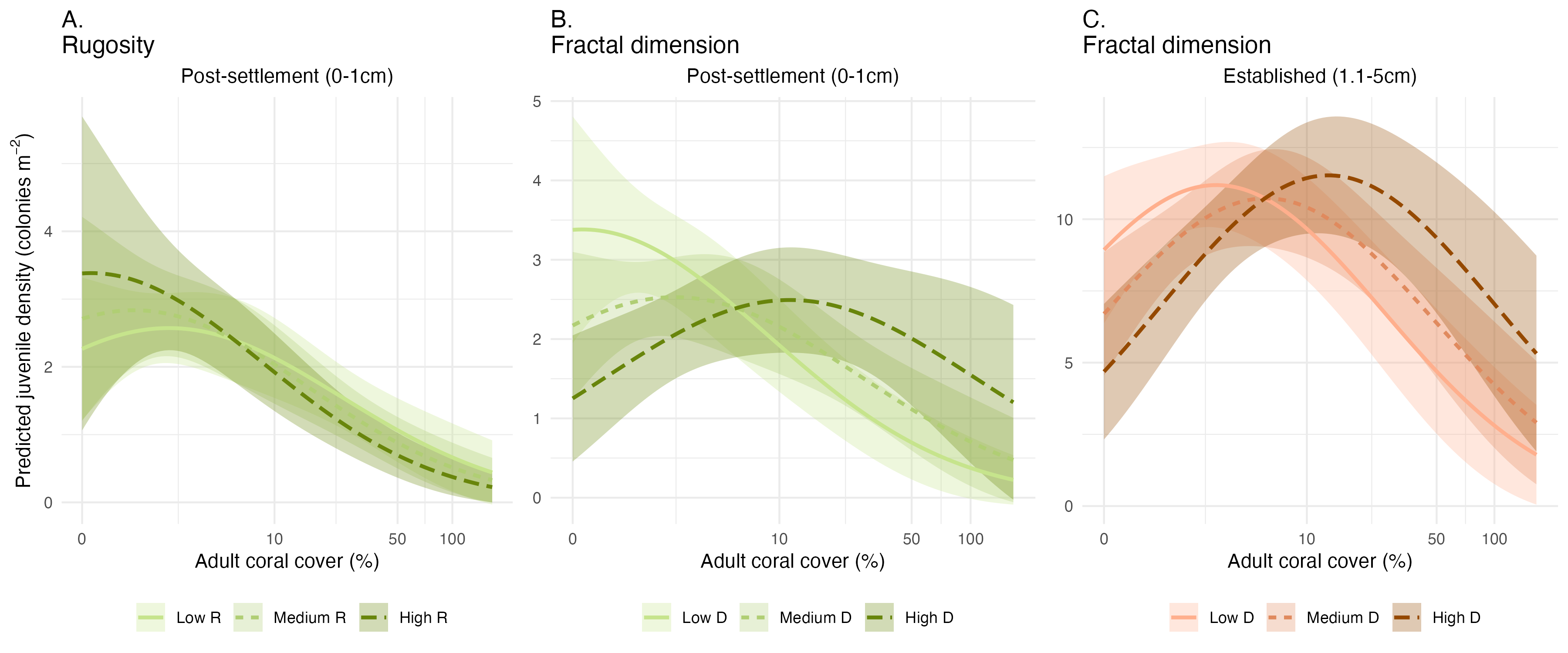

Figure S2. Significant effects of adult coral cover and reef structural complexity on two size classes of juveniles: post-settlement (0–1 cm) and established (1.1–5 cm). (A) Predicted post-settlement juvenile density as a function of adult coral cover and low, medium, and high rugosity. (B) Predicted post-settlement juvenile density as a function of adult coral cover and low, medium, and high fractal dimension. (C) Predicted established juvenile density as a function of adult coral cover and low, medium, and high fractal dimension. For all plots, shaded ribbons represent 95% confidence intervals and line types correspond to low, medium, or high levels of structure (10th, 50th, and 90th percentiles, respectively). Color corresponds to juvenile size class: post-settlement juveniles are green, established juveniles are orange. For predictions, all other covariates were held at their mean.

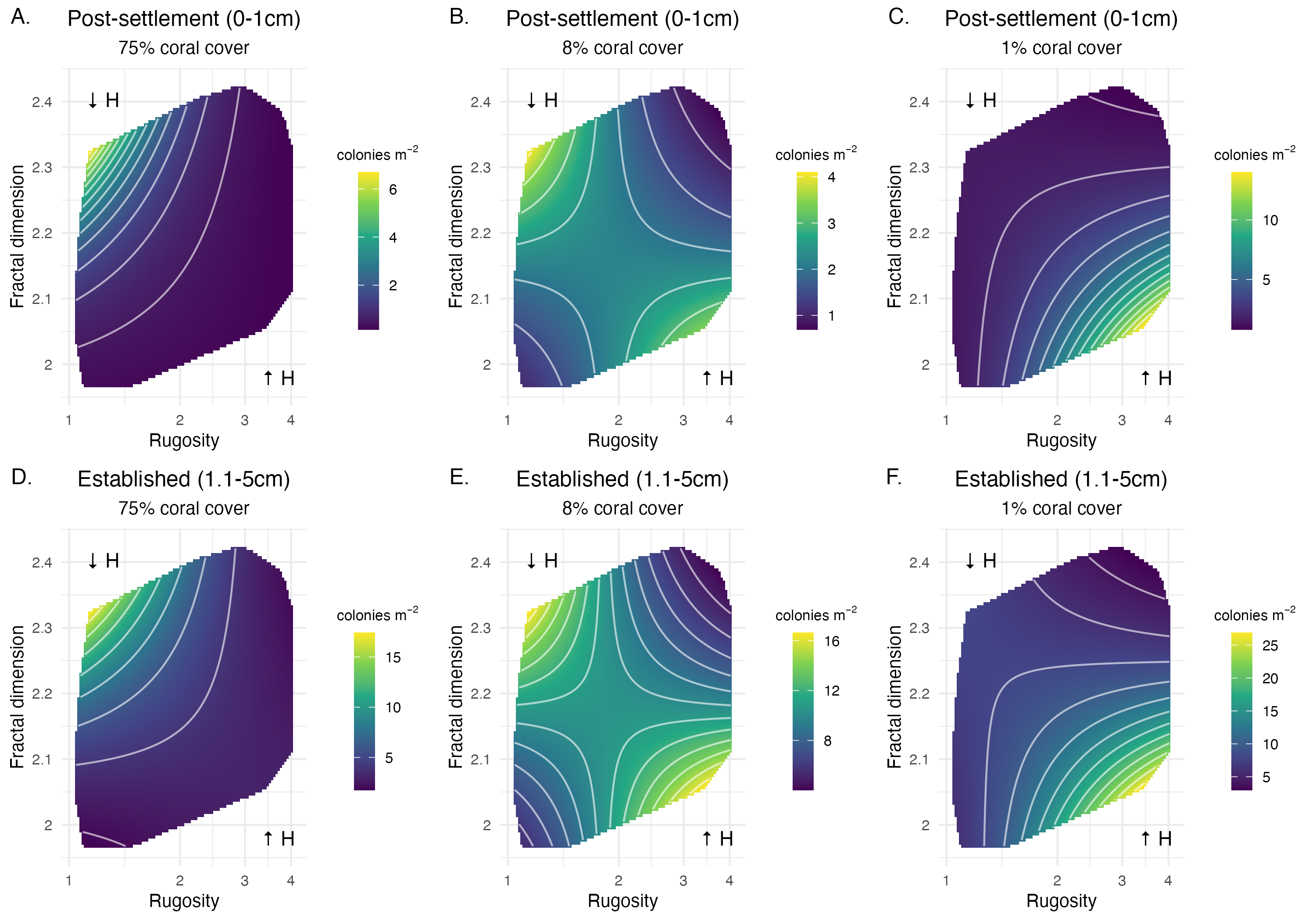

Figure S3. Predicted juvenile coral densities across combinations of rugosity and fractal dimension at fixed coral cover levels. (A-C) Predicted post-settlement (0–1 cm) juvenile densities and (D-F) predicted established (1.1–5 cm) juvenile densities at 75%, 8%, and 1% adult coral cover. Color scale shows predicted density and black points indicate observed juvenile densities. Arrows denote high (↑H) and low (↓H) height ranges according to the geometric relationship between rugosity and fractal dimension. All other covariates held at their mean.
